# Comparative analysis of MOB_Q4_ plasmids demonstrates that MOB_Q_ is a *cis*-acting-enriched relaxase protein family

**DOI:** 10.1101/726927

**Authors:** M. Pilar Garcillán-Barcia, Raquel Cuartas-Lanza, Ana Cuevas, Fernando de la Cruz

## Abstract

A group of small mobilizable plasmids is increasingly being reported in epidemiology surveys of enterobacteria. Some of them encode colicins, while others are cryptic. All of them encode a relaxase belonging to a previously non-described group of the MOB_Q_ class, MOB_Q4_. While highly similar in their mobilization module, two families with unrelated replicons can be distinguished, MOB_Q41_ and MOB_Q42_. Members of both groups were compatible between them and stably maintained in *E. coli*. MOB_Q4_ plasmids were mobilized by conjugation. They contain two transfer genes, *mobA* coding for the MOB_Q4_ relaxase and *mobC*, which was non-essential but enhanced the plasmid mobilization frequency. The origin of transfer was located between these two divergently transcribed *mob* genes. MPF_I_ conjugative plasmids were the most efficient helpers for MOB_Q4_ conjugative transmission. No interference in mobilization was observed when both MOB_Q41_ and MOB_Q42_ were present in the same donor cell. Remarkably, MOB_Q4_ relaxases exhibited a *cis*-acting preference for their *oriT*s, a feature already observed in other MOB_Q_ plasmids. These findings indicate that MOB_Q4_ plasmids can efficiently spread among enterobacteria aided by coresident IncI1, IncK and IncL/M plasmids, while ensuring their self-dissemination over highly-related elements.

**IMPORTANCE:** Plasmids are key vehicles of horizontal gene transfer and contribute greatly to bacterial genome plasticity. A group of plasmids, called mobilizable, is able to disseminate aided by helper conjugative plasmids. Here, we studied a group of phylogenetically-related mobilizable plasmids, MOB_Q4_, commonly found in clinically-relevant enterobacteria, uncovering the helper plasmids responsible for their dissemination. We found that the two plasmid species encompassed in the MOB_Q4_ group can coexist and transfer orthogonally, despite origin-of-transfer cross-recognition by their relaxases. Specific discrimination among their highly similar *oriT* sequences is guaranteed by the preferential *cis* activity of the MOB_Q4_ relaxases. Such strategy would be biologically relevant in a scenario of co-residence of non-divergent elements to favor self-dissemination.

## INTRODUCTION

Mobilizable plasmids are small genomes transmissible by conjugation. They encode a relaxase, and usually a relaxase accessory protein (RAP), which are in charge of the conjugative DNA processing at a specific site of the origin of transfer (*oriT*) called *nic*. Mobilizable plasmids lack the transfer genes required for establishing a conjugative bridge (mating pair formation system, MPF) to the recipient cell, as well as the type IV coupling protein (T4CP) that puts in contact relaxosome and MPF and thus depend on conjugative plasmids to be transferred (1).

According to their relaxase, transmissible plasmids were phylogenetically classified into MOB families (2) (3). Currently, nine relaxase MOB classes are defined, and five of them (MOB_P_, MOB_F_, MOB_Q_, MOB_H_ and MOB_C_) are prevalent in transmissible plasmids hosted in γ-Proteobacteria. Plasmids gathered in a relaxase MOB family share similar genomic traits. Relaxase MOB classification has thus shown to be a good predictor of the plasmid backbone (1, 4). Mobilizable plasmids resident in γ-Proteobacteria form phylogenetically-related clusters mainly within two relaxase MOB classes: MOB_P_ and MOB_Q_ (3). Relevant examples are ColE1-like plasmids, grouped in family MOB_P5_; IncQ1 plasmids, such as RSF1010/R1162, gathered in MOB_Q11_; and IncQ2 plasmids, such as pTC-F14, in family MOB_P14_ (1, 3). An additional clade of small plasmids encoding MOB_Q_ relaxases, previously classified as MOB_Qu_, and here redefined as MOB_Q4_, was observed in a phylogenetic reconstruction of this relaxase family (3).

A pair of degenerate primers specific for MOB_Q4_ plasmids was implemented in the Degenerate PCR MOB Typing (DPMT) approach developed by (5) to detect and classify transmissible plasmids. This method revealed the abundance of MOB_Q4_ plasmids in clinical isolates of enterobacteria (5, 6), previously unnoticed by other plasmid typing methods. Whole-genome sequencing of clinical *E. coli* isolates also uncovered the presence of this kind of plasmids (7-9). Prototype plasmids pIGWZ12 and ColE9-J (ColE2-like) cluster within the MOB_Q4_ clade. They are stable, theta-replicating, high copy-number, narrow host-range plasmids, whose replication systems have been extensively studied (10-14). Here, we uncovered the diversity of MOB_Q4_ plasmids, determined the helper conjugative plasmids responsible for their dissemination, and established their orthogonal behavior in terms of stability and transfer.

## MATERIALS AND METHODS

### Plasmid construction

MOB_Q4_ vectors were constructed by isothermal assembly of linear DNA fragments from PCR reactions, following the Gibson method (15, 16). The MOB_Q41_ backbone, based on the complete sequence of the pE2022_4 plasmid (GenBank Acc. No. KT693143 (8)), was linked to a kanamycin-resistance gene (coordinates 272 to 1216 of pSEVA211, GenBank Acc. No. JX560326 (17)) and a cerulean fluorescent protein gene (coordinates 41 to 1091 of pNS2-φVL (18)), producing plasmid pRC1 (see Supplementary Information S1). The MOB_Q42_ backbone was obtained by PCR amplification from the *E. coli* isolate HUMV 04/979 (6), which contains a ColE9-J-like plasmid (coordinates 1814 to 6114 of the sequence in Supplementary Information S2). It was joined to a chloramphenicol resistance gene (coordinates 272-1072 of pSEVA311, GenBank Acc. No. JX560331 (17)) and mCherry fluorescent protein gene (*cfp*, coordinates 1092-2117 of pNS2-φVL (18)), rendering plasmid pRC2 (see Supplementary Information S2). MOB_Q4_ vectors lacking the *mobC* ORF (from start to stop codon) were constructed by self-ligation of a single PCR fragment from either pRC1 or pRC2, generating plasmids pRC3 and pRC4, respectively.

Additional plasmids were constructed to delimit the *oriT* region. A schematic representation of the fragments included in each construction is depicted in Figure 1. Such fragments were individually assembled to coordinates 1-1030 and 1360-3001 of vector pSEVA631 (GenBank Acc. No. JX560348). Plasmids pRC5 and pRC6 contained a fragment including the *mobC* gene, the 178bp intergenic region between *mobC* and *mobA* and the first 400 nucleotides of the *mobA* gene from pRC1 and pRC2, respectively. Plasmids pRC7 and pRC8 included only the 178bp intergenic fragment (Supplementary Figure S1), located between genes *mobA* and *mobC* of pRC1 and pRC2, respectively. Plasmids pRC14 and pRC15 contain the *oriT* regions of pRC7 and pRC8 but cloned in the inverse orientation. Plasmids pRC11 and pRC9 respectively included portions 1-70 and 71-178 of the intergenic fragment between genes *mobA* and *mobC* of pRC1, while the same portions from pRC2 were included in pRC12 and pRC10, respectively. A pSEVA631 fragment containing coordinates 1-1030 and 1360-3001 was self-ligated, rendering the non-mobilizable vector pRC13, which was used as a control in the mating experiments.

**Figure 1.**
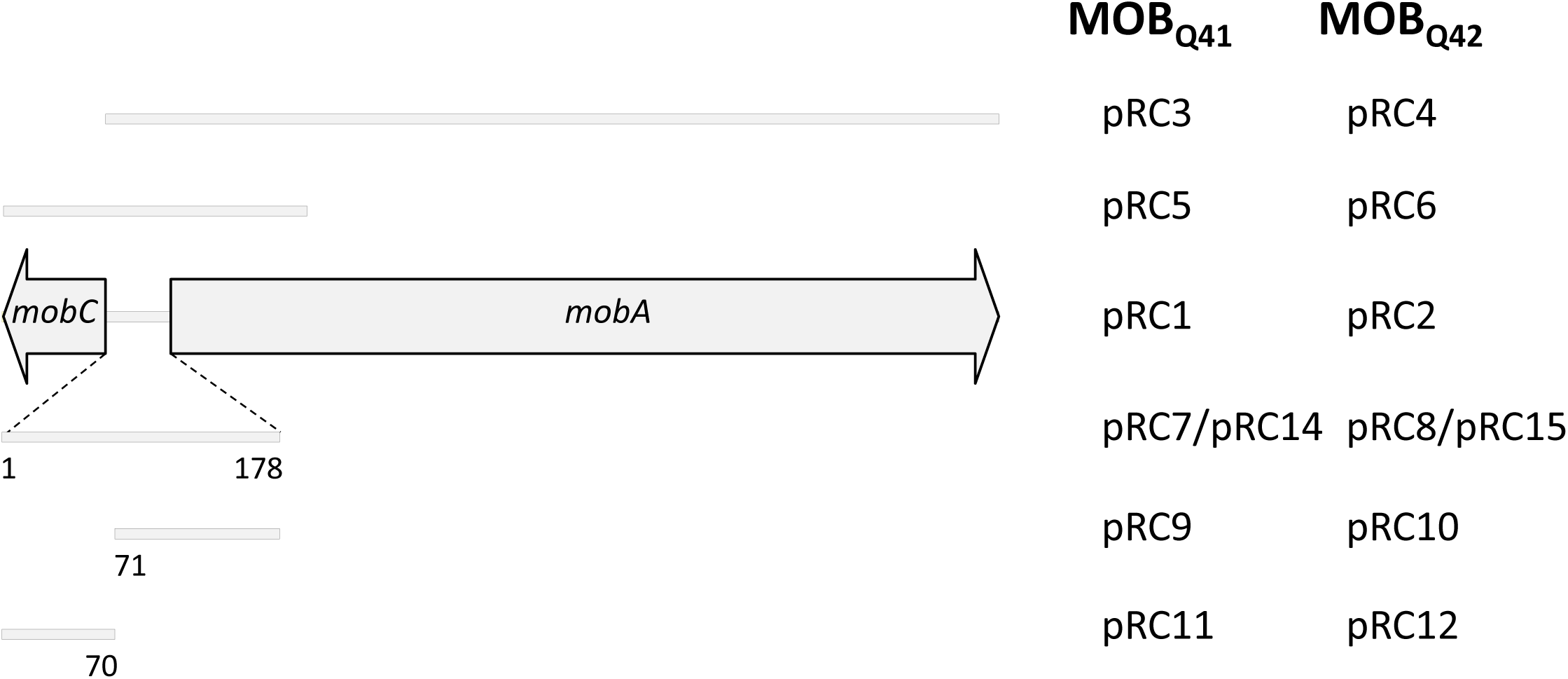
Schematic representation of the MOB_Q4_ DNA segments included in a series of recombinant plasmids. The mobilization region of MOB_Q4_ plasmids includes *mobC* and *mobA* genes, represented by large, horizontal gray arrows. The extent of the mobilization region included in each construction is represented by a gray bar. The plasmid names for the MOB_Q41_– based constructions are listed in the left column, while those for MOB_Q42_-based constructions are in the right column. Plasmids pRC1 and pRC2 also include the replication module of MOB_Q41_ and MOB_Q42_ plasmids, respectively.

### Stability assays

Plasmids pRC1 and pRC2 were independently introduced in the *recA*^+^ and *recA*^−^ isogenic strains UB1636 (19) and UB1637 (20) to check for their stability. Single colonies were inoculated in Lysogeny-Broth (LB) supplemented with kanamycin at 50 µg/ml (for pRC1-containing strains) or chloramphenicol at 25 µg/ml (for pRC2-containing strains) and grown to saturation at 37°C with agitation (150 rpm). A volume of 9.7 µl was transferred from saturated cultures to 10 mL of fresh LB media without antibiotics and grown to saturation in the same conditions. Rounds of transfer and growth were repeated up to 80 generations. The proportion of plasmid-bearing cells in the population was monitored by replica-plating 100 colonies in LB-agar supplemented with the appropriate antibiotics every 10 generations. A larger number of cells was inspected by fluorescence microscopy and, in the case of pRC1-containing cells, also by flow cytometry. Live cells were visualized using a Leica AF6500 microscope at 63x magnification. CFP and mCherry signals were monitored using BP filters (Excitation 434/17 – Emission 479/40 for CFP, Excitation 562/40 – Emission 641/75 for mCherry). Images were obtained using an iXon885 EM CCD Camera (Andor) and up to 1000 cells were analyzed in each case. Fluorescence emission was measured by flow cytometry using a FACS Canto II flow cytometer (Becton Dickinson) equipped with a 488 nm solid state laser for excitation. The cyan fluorescence of 20,000 events was detected using a 525/20 filter.

### Mating assays

Conjugative plasmids used in this work are listed in Supplementary Table S1. They were tested as helpers of MOB_Q4_ plasmids in surface mating experiments, following the procedure described by (21). Briefly, donor and recipient strains were mixed in a 1:1 ratio, deposited onto an LB-agar surface and incubated for 1h at 37°C (except when drR27 was used as a helper, in which case matings were carried out at 25°C). Then, the mixture was resuspended in LB and plated in the presence of appropriate antibiotics. Conjugation frequencies were expressed as the number of transconjugants per donor cell.

### Phylogenetic analysis

The 300 N-terminal residues of the MobA relaxase of plasmid ColE9-J were used as a query in a BLASTP search (22) (e-value: 1xE-3). The homologous sequences were aligned using MUSCLE (23). TrimAl v1.4 was used to calculate the average identity between sequences in the alignment (24). ProtTest 3 was used to estimate the best model of protein evolution for our set (25, 26). RAxML version 7.2.7 (27) was used for phylogenetic reconstruction. Using the JTTGAMMA model ten maximum likelihood (ML) searches trees were inferred and support values were assigned to each node of the best tree from 1000 bootstrap searches. Relaxase of the pXF5847 plasmid (GenBank Acc. no. YP_009076807.1) was used as outgroup.

### 3D structure prediction

Phyre2 was used to predict the 3D structure of the MobA relaxase domains of plasmids pE2022_4 and ColE9-J (28), which were visualized using PyMOL (29).

## RESULTS

### Analysis of MOB_Q4_ plasmids

The phylogenetic tree of MOB_Q4_ relaxases shows that they constitute a well-supported clade, which is in turn divided in two groups, MOB_Q41_ and MOB_Q42_ (Figure 2). The reconstruction was based on the first N-terminal 300 residues of MOB_Q4_ relaxases, which contain the three relaxase motifs (Figure 3A) and share 84% average amino acid identity (97 and 90% for individual MOB_Q41_ and MOB_Q42_ groups, respectively). Each MOB_Q4_ subclade groups highly related backbones (Figure 2). MOB_Q41_ are cryptic, small-size plasmids (Supplementary Table S2). Their backbone contains only four genes encoding a replication initiation protein (Rep), a relaxase (MobA), a putative relaxase accessory protein (MobC) and a hypothetical protein (Figure 2). The genes for the last two are generally not annotated. Besides the replication and mobilization genes, MOB_Q42_ plasmids also contain a colicin operon, including colicin, immunity and lysis genes, following the synteny of Group A nuclease colicins (30) (Figure 2). Plasmids ColE9-J and pO111_4 contain a second, partial colicin operon. The MOB_Q4_ subdivision in two relaxase groups matches with the presence of two different replicons (Supplementary Table S2). MOB_Q41_ plasmids encode a replication initiation protein that belongs to the Rep_3 superfamily (pfam PF01051), while MOB_Q42_ plasmids encode ColE2-like initiators (pfam PF03090 + PF08708). Thus, at least two plasmid species are included in MOB_Q4_, according to the criteria established by (31).

**Figure 2.**
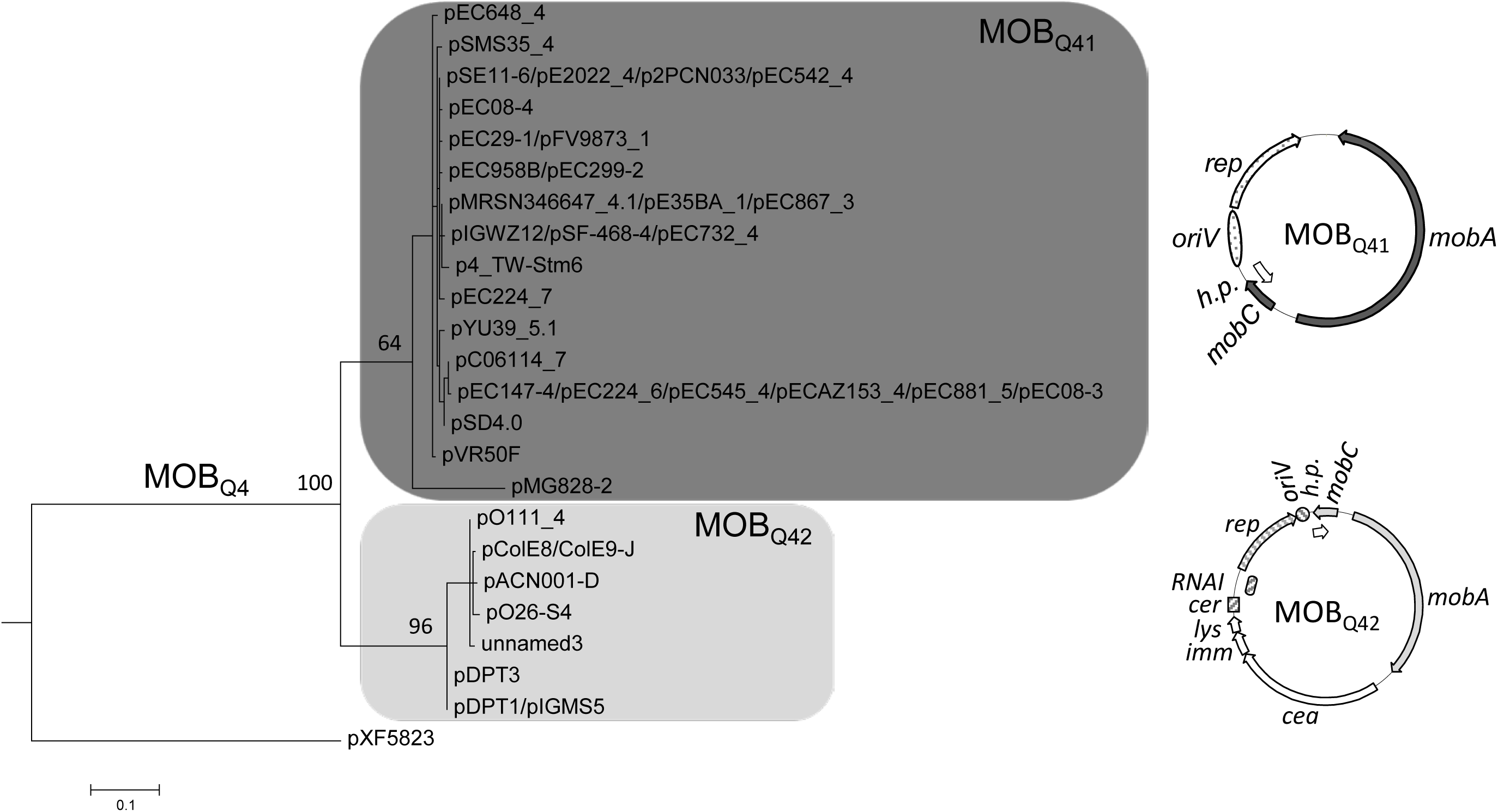
Phylogenetic tree of the MOB_Q4_ relaxase subclass. A maximum-likelihood phylogenetic reconstruction of the N-terminal domain of MOB_Q4_ relaxases is shown. MobA relaxase of plasmid pXF5843 were used as outgroup. Bootstrap values of relevant nodes are indicated. Families Q41 and Q42 are shadowed in dark and light gray, respectively. A prototype backbone of each MOB_Q4_ subfamily is represented to the right of the corresponding clade. The common genes of the mobilization module are represented in the same gray color pattern. The elements of the replication module are dotted (for MOB_Q41_) or striped (for MOB_Q42_). Genes of the colicin operon are depicted in a white background.

**Figure 3.**
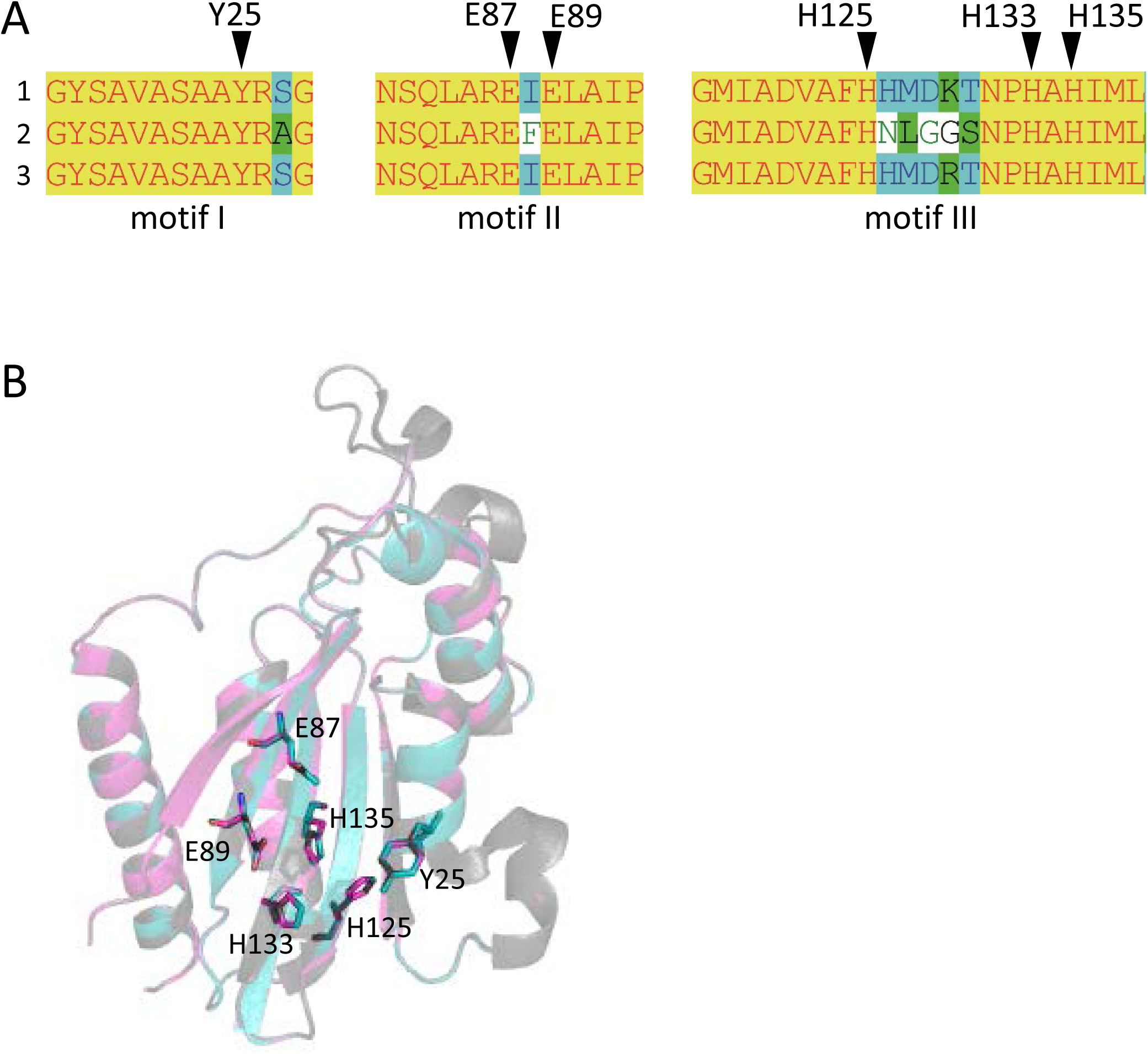
The MOB_Q4_ relaxase domain. A) Multiple alignment of the three conserved MOB_Q_ relaxase motifs (3). (1) MOB_Q41_ relaxases, with the exception of pMG828-2; (2) pMG828-2; (3) MOB_Q42_ relaxases. Putative catalytic residues are indicated by black triangles over the amino acid sequences. B) Protein 3D-structure superposition of MOB_Q41_, MOB_Q42_ and MOB_Q11_ relaxases. Structure prediction of MOB_Q4_ relaxases was carried out with Phyre2 (28). In gray, the minimal relaxase domain of MobA of plasmid R1162 / RSF1010 used as a model (PDB 2NS6); in blue and magenta the predicted structures of the homologous domains of the MOB_Q41_ (MobA of pE2022_4) and MOB_Q42_ (MobA of ColE9-J) relaxases, respectively. Key residues of MOB_Q4_ relaxases are highlighted as sticks.

The 3D structure prediction of the relaxase domain of MOB_Q41_ and MOB_Q42_ plasmids pE2022_4 and ColE9-J rendered MOB_Q_ relaxases NES (plasmid pLW1043, PDB Acc. No. 4HT4, (32)) and MobA (plasmid R1162 / RSF1010, PDB Acc. No. 2NS6, (33)) as best hits (100% confidence). The superimposed structures pointed to MOB_Q4_ amino acids Y25 (motif I), E87 and E89 (motif II), and H125, H133 and H135 (motif III) as homologs of the MobA_R1162 catalytic residues Y25, E74 and E76, and H112, H120 and H122, respectively (Figure 3B). Contrary to the high conservation of the N-terminal domain among members of both MOB_Q4_ subgroups, the amino acid identity of the C-terminal part of the MOB_Q4_ relaxases dropped to 35%. This C-terminal domain exhibited low homology to SogL primases of IncI1 plasmids.

### Stability and co-residence of MOB_Q4_ plasmids

To study the MOB_Q4_ plasmids, two synthetic plasmids were constructed, pRC1 and pRC2. They included the replication and mobilization modules of MOB_Q41_ and MOB_Q42_ backbones, respectively. Antibiotic-resistance (AbR) and fluorescent protein genes were also included as reporters for the stability and mating experiments. Plasmid stability was assayed in *recA*^+^ and *recA*^−^ *E. coli* strains, containing either one or both plasmids (to test for stability and incompatibility, respectively). In all cases, the percentage of plasmid retention in the bacterial population after 80 generations was 100%. Thus, besides stability in *E. coli*, both MOB_Q4_ plasmid species exhibited full compatibility.

### Mobilization of MOB_Q4_ plasmids by different MPF systems

Conjugative plasmids encoding three different MPF systems, MPF_T_, MPF_I_ and MPF_F_, the most abundant in γ-Proteobacteria, were used as helpers to mobilize the MOB**_Q4_** plasmids pRC1 and pRC2 (Table 1 and Figure 4). Their mobilization by MPF_F_ plasmids (pOX38, R100-1 and drR27) was not detected. Three of the MPF_T_-type helper plasmids (R388, pKM101, and pOLA52) did not mobilized pRC1 neither pRC2, while pRL443, R751, and R6K*drd1* did mobilize them. pRL443 and R751 are closely related plasmids, respectively belonging to the α and β subdivisions of the IncP1 group. Nevertheless, their efficiency at mobilizing MOB_Q4_ plasmids differed widely, being that of IncP1α plasmid approximately 1000-fold higher. Both MPF_I_ plasmids (R64*drd11* and pCTX-M3) mobilized MOB_Q4_ plasmids efficiently. The highest mobilization frequencies obtained for both MOB_Q4_ plasmids, pRC1 and pRC2, were achieved by using R64*drd11* as a helper. Besides, the mobilization efficiency of the MOB**_Q4_** plasmids in relation to the transfer of the helper conjugative plasmid was also the highest for R64*drd11* (Figure 5). So, R64*drd11* was the most efficient helper plasmid.

**Table 1.** Mobilization frequencies of MOB_Q4_ plasmids by different MPF type helpers.

| Mobilizable plasmid | MPF type of the helper plasmid |  |  |  |  |  |  |  |  |  |  |
| --- | --- | --- | --- | --- | --- | --- | --- | --- | --- | --- | --- |
|  | MPF<sub>T</sub> |  |  |  |  |  | MPF<sub>I</sub> |  | MPF<sub>F</sub> |  |  |
|  | <u>RP4</u> | <u>R751</u> | <u>R388</u> | <u>pKM101</u> | <u>R6Kdrd1</u> | <u>pOLA52</u> | <u>R64drd11</u> | <u>pCTX-M3</u> | <u>pOX38</u> | <u>R100-1</u> | <u>R27dr</u> |
| pRC1 (MOB<sub>Q41</sub>) | 1.6x10<sup>-2</sup><br>(1x10<sup>-2</sup> – 2.7x10<sup>-2</sup>) | 1.7x10<sup>-4</sup><br>(2.1x10<sup>-5</sup> – 7.6x10<sup>-5</sup>) | <1x10<sup>-7</sup> | <1x10<sup>-7</sup> | 1.1x10<sup>-6</sup><br>(2.7x10<sup>-7</sup> – 4.3 x10<sup>-6</sup>) | <1x10<sup>-7</sup> | 5.1x10<sup>-1</sup><br>(3.9x10<sup>-1</sup> – 6.7x10<sup>-1</sup>) | n.d. | <1x10<sup>-7</sup> | n.d. | <1x10<sup>-7</sup> |
| pRC2 (MOB<sub>Q42</sub>) | 1.2x10<sup>-1</sup><br>(8.4x10<sup>-2</sup> – 1.8x10<sup>-1</sup>) | 7.8x10<sup>-5</sup><br>(6.3x10<sup>-5</sup> – 9.7x10<sup>-5</sup>) | <1x10<sup>-7</sup> | <1x10<sup>-7</sup> | 1.1x10<sup>-4</sup><br>(7.8x10<sup>-5</sup> – 1.6 x10<sup>-4</sup>) | <1x10<sup>-7</sup> | 1.2<br>(4.6x10<sup>-1</sup> – 3.2) | 1.0x10<sup>-3</sup><br>(4.5x10<sup>-4</sup> – 2.4x10<sup>-3</sup>) | <1x10<sup>-7</sup> | <1x10<sup>-7</sup> | <1x10<sup>-7</sup> |

n.d.: Not determined due to the lack of an appropriate AbR marker.

**Figure 4.**
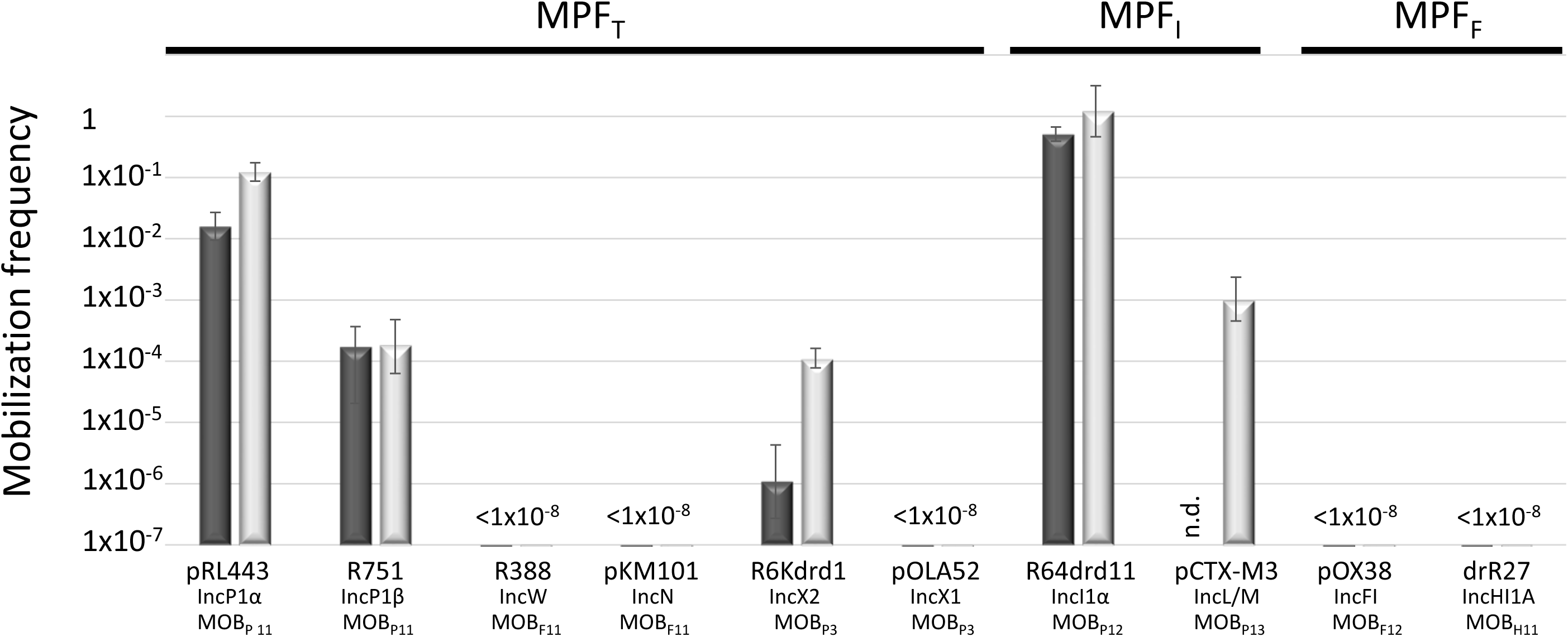
Mobilization frequencies of MOB_Q4_ plasmids by a series of helper plasmids. The mobilization frequency was calculated as the number of transconjugants containing the MOB_Q4_ plasmid per donor cell. Figures are the average of at least six independent experiments. Dark- and light-gray bars indicate the mobilization frequencies of pRC1 (MOB_Q41_) and pRC2 (MOB_Q42_) plasmids, respectively. Below the bars the helper plasmid used in each case, as well as its corresponding Inc and MOB groups, are indicated. The MPF types of the helper plasmids are indicated in the upper part of the figure.

**Figure 5:**
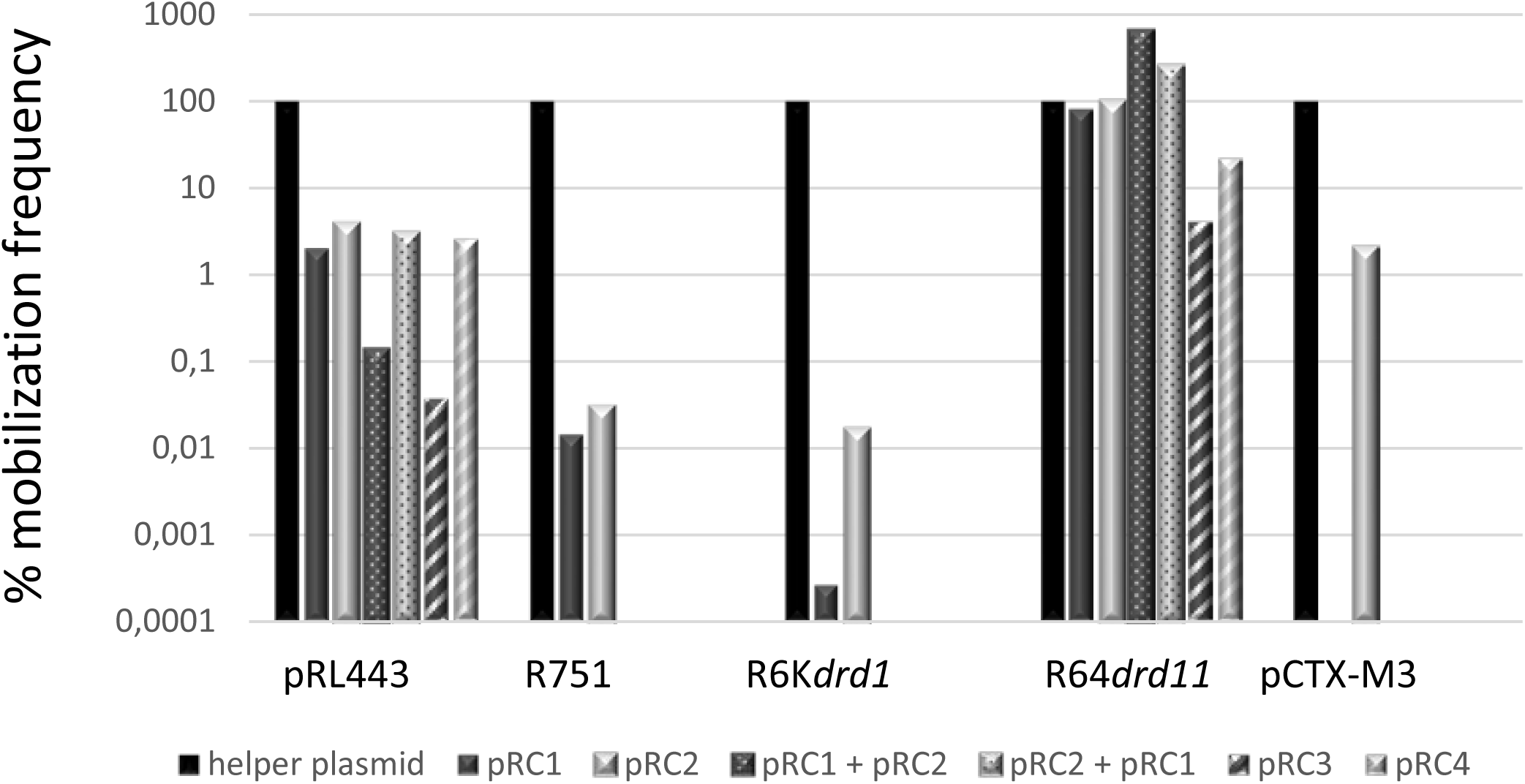
Mobilization frequencies of different plasmid combinations. The mobilization efficiencies of the MOB_Q4_ plasmids are relativized to the helper plasmid transfer rates (black bars, 100%). pRC1 and pRC2 efficiencies are represented by dark- and light-gray bars, respectively. pRC1 + pRC2 (dotted bar in a dark-gray background) indicates the mobilization frequency of pRC1 when coresident with pRC2. pRC2 + pRC1 (dotted bar in a light gray background) indicates the mobilization frequency of pRC2 when coresident with pRC1. The mobilization efficiencies of the *mobC*^−^derivatives pRC3 and pRC4 are represented by striped bars in dark and light gray backgrounds, respectively. The bars represent the average of at least six experiments.

To test whether the mobilization of the MOB_Q41_ plasmid was affected by co-residence with a MOB_Q42_ plasmid and vice versa, pRC1 and pRC2 were introduced conjointly with the helper plasmid (either pRL443 or R64*drd11*) in the same host. The values obtained for pRC1 and pRC2 mobilization when residing together in the same host did not differ significantly from those obtained when only one of them was present in the donor cells (Figure 5).

### *mobC* deletion effect in the mobilization efficiency

To check whether MobC plays a role in the MOB_Q4_ plasmid mobilization, *mobC* deletion mutants were constructed from pRC1 and pRC2, respectively producing pRC3 and pRC4 (Figure 1). A moderate decrease in mobilization was observed in the *mobC*^−^variants (two-log reduction for pRC3 and one-log for pRC4, when using R64*drd11* as a helper (Figure 5)).

### *In trans* mobilization of *oriT*_MOB_Q4_-containing vectors

The 178 bp intergenic region comprised between the *mobC* and *mobA* genes of MOB_Q4_ plasmids was assembled with an *oriT*-lacking fragment of vector pSEVA631. The resulting constructions, pRC7 (for MOB_Q41_) and pRC8 (for MOB_Q42_) (Figure 1), were introduced in donor strains to check for their mobilization. The transfer proteins were supplied *in trans*: the corresponding mobilizable plasmid (pRC1 or pRC2) provided the relaxosomal proteins, while the conjugative plasmid (R64*drd11*) supplied the T4CP and MPF. Plasmids pRC7 and pRC8 were transferred to the recipient population, but 1000-fold less efficiently than their corresponding *mobA*^+^*mobC*^+^ partners (pRC1 and pRC2) (Figure 6). This result was confirmed by using plasmids pRC14 and pRC15, instead of pRC7 and pRC8, in the mobilization experiments. Plasmids pRC14 and pRC15 contained the same *oriT* region present in pRC7 and pRC8, but cloned in the inverse orientation. Besides, to avoid losing any *oriT*-related function, larger segments including also the *mobC* gene and the first 431bp of the *mobA* gene (pRC5 and pRC6 (Figure 1)), were analyzed. Here again relaxase, T4CP and MPF components were provided *in trans*. Plasmids pRC5 and pRC6 behave similarly to pRC7 and pRC8, and were mobilized at least 500-fold less than pRC1 and pRC2 (Figure 6).

**Figure 6:**
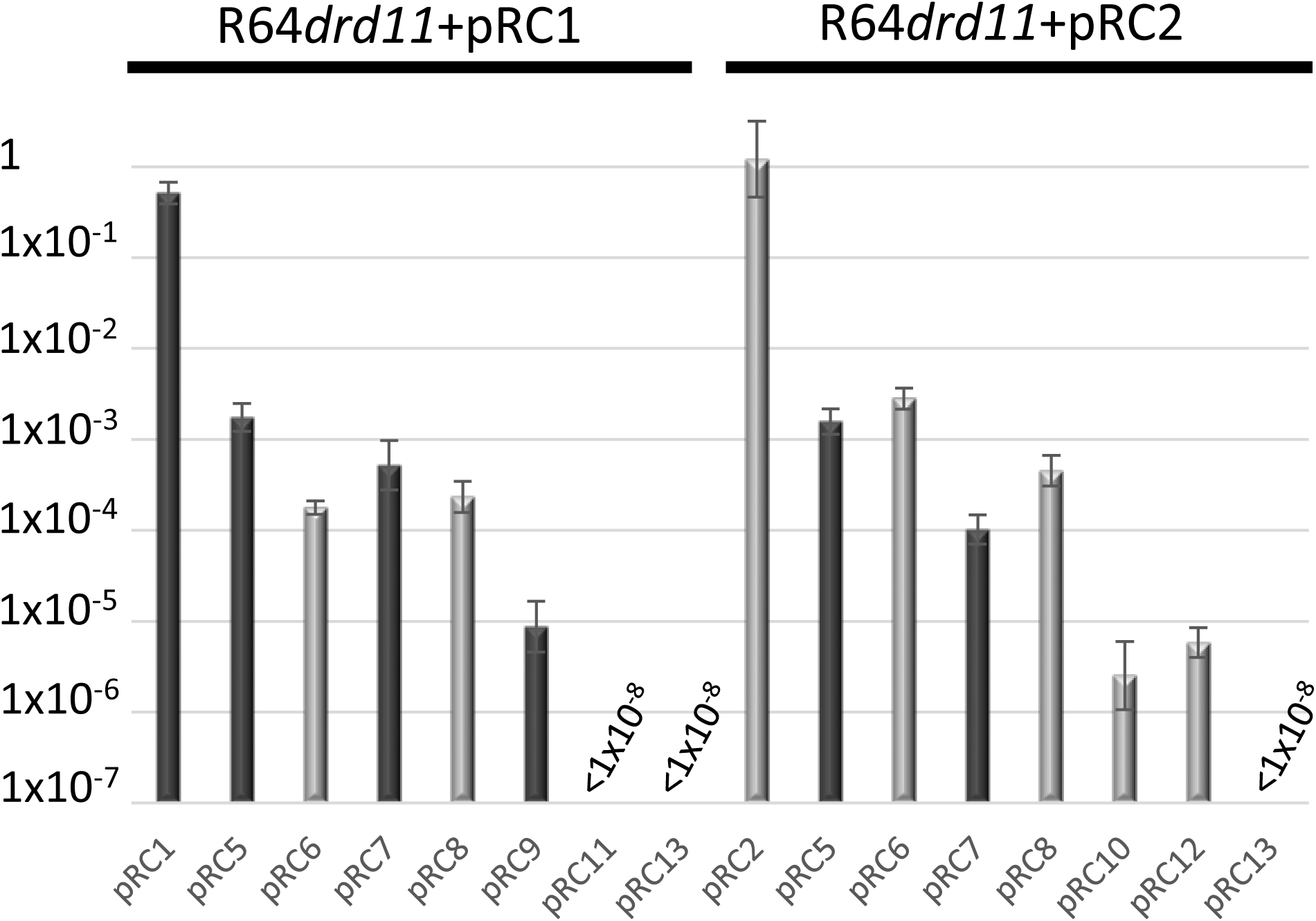
In *trans* mobilization of *oriT* fragments. R64*drd11*-mediated mobilization frequencies of pRC1 and plasmids containing fragments of its *oriT* are represented by dark-gray bars, while those of pRC2 and its derivatives are in light-gray bars. The bars represent the average of at least six experiments.

To further delimit the *oriT* of MOB_Q4_ plasmids, the 178bp *oriT* fragments cloned in pRC7 and pRC8 (see Supplementary Figure S1) were subdivided in two portions, one containing *oriT* nucleotides 1-70 (pRC11 and pRC12) and the other containing *oriT* nucleotides 71-178 (pRC9 and pRC10) (Figure 1 and Supplementary Figure S1). Disruption of the 178bp *oriT* region resulted in a drastic loss of conjugation efficiency (Figure 6).

The MOB_Q4_ relaxases were also tested for their specificity to act on a non-cognate MOB_Q4_ *oriT*. Mobilization frequencies of *oriT*_MOB_Q42_ plasmids pRC6 or pRC8 by the MOB_Q41_ plasmid pRC1 + R64*drd11*, as well as *oriT*_MOB_Q41_ plasmids pRC5 or pRC7 by the MOB_Q42_ plasmid pRC2 + R64*drd11*, were similar to that obtained for the cognate systems (Figure 6).

## DISCUSSION

MOB_Q_ is a broad relaxase class that encompasses several families, each of which includes related plasmid backbones: MOB_Q1_ comprises relaxases of mobilizable broad host-range IncQ1-like plasmids; MOB_Q2_, conjugative relaxases of pTi and many rhizobial plasmids; MOB_Q3_, conjugative broad host-range plasmids resident in gram-positive, such as pIP501 (3). In the referred study, many MOB_Q_ plasmids were not ascribed to a specific subclassification due to either low resolution of the clades or lack of information on the plasmid members. Here, we focused on one of these poorly defined clades, now named MOB_Q4_, prompted by the fact that these relaxases have been recurrently detected in enterobacterial clinical isolates (5-9).

MOB_Q4_ relaxases formed two clusters that correlated with the presence of two different replicons, thus containing at least two plasmid species as defined by (31) (Figure 2 and Supplementary Table S2). Plasmids pIGWZ12 and ColE9-J exemplify each cluster. They are stable, theta-replicating, high copy number plasmids (15 and 10 copies per chromosome molecule, respectively (34, 35)). The origin of replication of plasmid pIGWZ12 was located upstream the *rep* gene. It contains iterons, an A+T rich region and four DnaA boxes (10, 11). The iterons were found to be the incompatibility determinants (10). The corresponding replication initiation protein belongs to the Rep_3 superfamily (PF01051) (11). ColE2-like plasmids, such as ColE9-J, form a group of closely related elements that share an identical priming mechanism, mediated by the plasmid-encoded Rep protein (13, 36-38). The origin of replication consists of 32 bp located downstream of the *rep* gene, containing two directly repeated sequences (12, 39-41). In ColE2-like plasmids, the *rep* gene expression is post-transcriptionally controlled by a plasmid-encoded RNA (*RNAI*), which binds the untranslated 5’ region of the *rep* mRNA, preventing its translation (14, 35, 42). MOB_Q42_ plasmids contain a *cer*-like site (38), an indication that they profit from a host site-specific recombination system for resolving multimers to monomers as ColE1-like plasmids do (43-45).

Completely sequenced MOB_Q4_ plasmids all originate in Enterobacteriaceae family hosts (Supplementary Table S2). They were isolated from different backgrounds: *Salmonella enterica* isolated from pork meat (pSD4.0) (46), pork feces (p4_TW-Stm6) (47) and human systemic infection (pYU39_5.1) (48), multidrug-resistant environmental *E. coli* (pSMS35_4) (49), commensal *E. coli* (pSE11-6) (50), enterohemorrhagic *E. coli* strains of the O26 and O111 serogroups (pO26-S4 and pO111_4) (51, 52), extended-spectrum beta-lactamase producing *E. coli* clinical isolates (pE2022_4, pFV9873_1, pEC147-3 and pEC08-6) (7, 8), *E. coli* isolated from human urinary tract (pVR50F) (53) and bloodstream infections (pSF-468-4) (54), as well as porcine extraintestinal pathogenic *E. coli* strain (p2PCN033) (55), among others. None of these plasmids contain AbR genes. There is still no clue on the selective advantage provided by the cryptic MOB_Q41_ plasmids. In the case of MOB_Q42_ plasmids, the fact that all carry colicin operons, a priori an advantageous trait for the bacterial host, could explain the abundance of this type of plasmids. For example, the MOB_Q42_ plasmid pDPT1 was stably acquired by a Vietnamese *Shigella sonnei* strain in the mid-1990s, and became fixed in the evolving bacterial population (56). The colicin E5 produced by pDPT1 was highly bactericidal against nonimmune *Shigella* and *E. coli* strains. The acquisition of the pDPT1 colicin plasmid, coinciding with the high increase of dysentery produced by this strain, suggests that pDPT1 conferred a beneficial function to its host (56).

To get insight into the MOB_Q4_ family, mobilizable synthetic plasmids based on two prototype backbones (pE2022_4 from MOB_Q41_ and ColE9-J from MOB_Q42_) were constructed (pRC1 and pRC2). They included the natural replication and mobilization modules, and AbR and fluorescent protein genes as markers. Firstly, they were assayed for their maintenance in the bacterial population. Stability experiments indicated that MOB_Q4_ backbones were highly stable in *E. coli* (100% retention in 80 generations) despite the cargoes loaded in plasmids pRC1 and pRC2. Furthermore, both plasmids were 100% stable when replicating in the same host cell, which indicates that they are compatible and do not interfere with the stable vertical inheritance of each other.

Since mobilizable plasmids do not encode the mating pair formation system neither the T4CP, their transfer relies on auto-transmissible plasmids. We wondered which conjugative plasmids could be responsible for the dissemination of MOB_Q4_ plasmids. Not all conjugative plasmids are equally efficient at supplying these functions to a specific mobilizable plasmid (57, 58). The contacts established between the relaxosome of the mobilizable plasmid and the T4CP-MPF of the helper plasmid are crucial in the transfer process. ColE1-like MOB_P5_ plasmids are efficiently mobilized by IncF-MOB_F12_ (*e.g.* F) and IncI1-MOB_P12_ (*e.g.* R64*drd11*) plasmids (58). IncQ1-MOB_Q1_ plasmids, such as RSF1010, are transferred by IncP1-MOB_P11_ helper plasmids (*e.g.* RP4) (58, 59). pMV158-like plasmids (MOB_V1_) are mobilized by IncP1-MOB_P11_ and Inc18-MOB_Q3_ (*e.g.* pIP501) plasmids (60).

We looked for reports providing indirect evidence on MOB_Q4_ plasmid mobilization through conjugation. In a survey for the presence of transmissible plasmids in a multidrug *E. coli* collection, MOB_Q4_ transconjugants were obtained from seven out of the eight MOB_Q4_ containing clinical isolates (6). In all cases, a MOB_P12_-MPF_I_ plasmid, presumptively the helper, was also present in both, donor and transconjugant cells. Similarly, the MOB_Q41_ plasmid pSD4.0 and the IncI1 plasmid pSD107 were found in *E. coli* transconjugants arisen from a mating with *Salmonella enterica* (46).

Three conjugative MPF types (MPF_T_, MPF_F_, and MPF_I_) are prevalent in Enterobacteriaceae (61, 62), the taxonomic class where MOB_Q4_ plasmids have been found. In this study, a set of conjugative plasmids representative of these MPF families were tested as helpers for the mobilization of MOB_Q4_ plasmids (Figures 4 and 5). Not all MPF types were equally efficient. R64*drd11*, the prototype of IncI1α-MOB_P12_ plasmids, which encodes a MPF_I_ conjugative apparatus, was the most efficient helper. Plasmid pCTX-M3 (IncL/M-MOB_P13_), was also an efficient helper. Co-residence of MOB_Q4_ with MPF_I_ plasmids has been observed for MOB_Q4_ plasmids pSE11-6 (50), pSD4.0 (46), pEC147-4 (7), pO26-S4 (Fratamico *et al.*, 2011), pDPT1 (56) and pE2022_4 (8).

On the other hand, MPF_F_-type plasmids (e.g. IncF-MOB_F12_ (F) or IncHI1-MOB_H11_ (R27) plasmids), which show high prevalence in enterobacteria, were not appropriate for MOB_Q4_ mobilization. MPF_T_ plasmids behaved unevenly as MOB_Q4_ mobilizers. IncP1-MOB_P11_ (RP4 and R751) and IncX2-MOB_P3_ (R6K*drd1*) plasmids rendered MOB_Q4_ transconjugants, while IncW-MOB_F11_ (R388), IncN-MOB_F11_ (pKM101) or IncX1-MOB_P3_ (pOLA52) did not. Contrary to IncP, IncW and IncN plasmids, most IncF, IncI1, IncH and IncX plasmids are naturally repressed for conjugation. In this study, we used derepressed variants of IncF (pOX38 and R100-1), IncI1α (R64*drd11*), IncHI1 (drR27) and IncX2 (R6K*drd1*) plasmids, but not a derepressed IncX1. IncX1 and IncX2 plasmids are highly similar in their conjugation genes. Taking into account that the IncX2 derepressed plasmid R6K*drd1* was not efficient at mobilizing MOB_Q4_ plasmids (Table 1, Figures 4 and 5), and that the IncX1 plasmid pOLA52 self-transfers at low frequency (around 10^−4^ per donor)(63), the lack of mobilization of the MOB_Q4_ plasmids pRC1 and pRC2 by pOLA52 is not surprising. The widely different mobilization efficiencies displayed by the two IncP1 helpers used is more curious. RP4 and R751 are prototypes of the α and β divisions of IncP1 backbones, respectively. Despite the high conservation of their transfer genes, the kanamycin-sensitive RP4 derivative, pRL443, was 100-1000 times more efficient than R751 as a MOB_Q4_ helper. Noticeable differences were observed for these two conjugative plasmids at transferring IncQ2-MOB_P14_ mobilizable plasmids pTC-F14 and pTF-FC2 (64).

Many conjugative and mobilizable plasmids encode RAPs that recognize and bind their cognate *oriT* sequence probably favoring a single-stranded state around the *nic* site (65). Deletion of RAP genes *trwA* of R388 (66), *nikA* of R64 (67), *mobB* and *mobC* of plasmids pTC-F14 and pTF-FC2 (64), *traJ* and *traK* of RP4 (68), *mobC* of R1162/RSF1010 (69), and *mbeC* of ColE1 (70) resulted in drastic decrease of plasmid transfer. All MOB_Q4_ plasmids encode a gene, called *mobC*, which is located adjacent to *oriT* and transcribed opposite to the *mobA* relaxase gene (Figure 1). Most of the *mobC* genes are not annotated, so we updated their annotation, as listed in Supplementary Table 2. The MobC proteins of MOB_Q4_ plasmids are small (less than 100 amino acids) and showed no homology to other RAPs (by using PSI-Blast). Here, we demonstrated that they are not absolutely essential for MOB_Q4_ plasmid mobilization. Nevertheless, *mobC* deletion resulted in 10 to 100-fold reduction in MOB_Q4_ plasmid mobilization frequencies (Figure 5). This is an interesting difference to other plasmid groups, which should be further investigated. It is conceivable that some MOB_Q4_ plasmids can be found, the mobilization of which is independent of RAPs.

We delimited the *oriT* of MOB_Q4_ plasmids to the 178 bp intergenic region between genes *mobC* and *mobA* (Supplementary Figure S1). *oriT*-containing plasmids (pRC5 and pRC7 for MOB_Q41_, or pR6 and pRC8 for MOB_Q42_) were mobilized by the relaxase proteins provided by the corresponding mobilizable plasmid (either pRC1 or pRC2) and the T4CP+MPF functions of the helper plasmid (R64*drd11*) (Figure 6). Disruption of such 178 bp segment resulted in a drastic loss of conjugation efficiency of the *oriT-*containing plasmid, as previously reported for pIGWZ12 (10). Furthermore, MOB_Q41_ relaxases were able to recognize and process the heterologous *oriT*_MOB_Q42_ with the same efficiency as the cognate *oriT*_MOB_Q41_. The same situation accounted for MOB_Q42_ relaxases (Figure 6). These two *oriT*s showed 10 differences in their 178bp sequence.

Remarkably, MOB_Q4_ relaxases showed a *cis*-acting preference for their *oriT*s. They acted at least 500-fold better on a *cis* than on a *trans oriT* substrate (Figure 6). The *cis-*acting preference is a characteristic exhibited by some DNA-binding proteins, such as the TnpA transposases of Tn*10*, Tn*5* and Tn*903* (71-73). Relaxases generally lack a *cis* preference for their *oriT*s. There are only a few examples of relaxases that show preference for a *cis*-encoded substrate. Notably, all plasmid-encoded *cis*-acting relaxases have been reported in members of the MOB_Q_ class: TraA of plasmid pRetCFN42d (MOB_Q2_) (74) and TraA of plasmid pIP501 (MOB_Q3_) (75). The MOB_P_ relaxase of transposon Tn*1549* was also found to be *cis*-acting (76). Nevertheless, other MOB_Q_ relaxases, such as Nes_pSK41 (77), as well as MobA of plasmids R1162 / RSF1010 and pSC101 (69, 78, 79) worked efficiently in *trans*. The *cis*-acting preference of the MOB_Q4_ relaxases implies that when two MOB_Q4_ plasmids are present in the same cell, the contribution of *oriT* cross-recognition by the heterologous MOB_Q4_ relaxase to plasmid transfer is not substantial. This feature could be essential to guarantee their efficient transfer, given the fact that both types of MOB_Q4_ plasmids use the same repertoire of conjugative helpers.

## CONCLUSIONS

MOB_Q41_ and MOB_Q42_ plasmids are orthogonal in terms of replication and mobilization. That is, they are able to coexist stably in the *E. coli* population without negatively interfering between each other. They disseminate through bacterial conjugation, aided specially by MPF_I_ conjugative plasmids, but neither of the MOB_Q4_ plasmids dominates the horizontal transfer process. Co-residence of MOB_Q41_ and MOB_Q42_ plasmids in the same host neither hindered nor boosted their respective mobilization frequencies. So, MOB_Q41_ relaxosome interaction with the transfer machinery of the helper plasmid was not affected by the presence of a MOB_Q42_ plasmid and vice versa. Since both plasmids (MOB_Q41_ and MOB_Q42_) circulate among enterobacteria, their coexistence in natural environments is likely. In such ecological setting, specific discrimination among their highly similar *oriT* sequences would be guaranteed by the preferential *cis* activity of the MOB_Q4_ relaxase. Such strategy would be biologically relevant in a scenario of co-residence of non-divergent elements to favor self-dissemination.

## ACKNOWLEDGMENTS

This work was supported by the Spanish Ministry of Economy and Competitiveness (BFU2017-86378-P to FC) and Consejo Superior de Investigaciones Científicas (201820I143 to MPGB). The authors want to thank María Aramburu and Raúl Fernández-López for their technical assistance with the flow cytometer and the fluorescence microscopy, respectively.

